# Secretome analysis of cancer-associated fibroblasts from prostate cancer to identify potential therapeutic targets

**DOI:** 10.1101/2024.12.09.627457

**Authors:** Apoorva Abikar, Mohammad Mehaboob Subhani Mustafa, Radhika Rajiv Athalye, Namratha Nadig, Ninad Tamboli, Vinod Babu, Ramaiah Keshavamurthy, Prathibha Ranganathan

## Abstract

Tumor microenvironment (TME) is a complex entity comprising of several cell types secreted factors as well as an extracellular matrix. A dynamic interaction between tumor cells and their environment profoundly influences tumor survival, aggressiveness, and progression. Cancer-associated fibroblasts (CAFs) are one of the major cellular components of TME and serve as a major source of various secreted factors. These factors are known to modulate tumor survival and progression, as well as their response to therapy. Despite the importance of the TME factors on various aspects of tumor cell behavior, to date factors unique to CAFs that could be potential therapeutic targets are not identified in most systems. This study was aimed at identifying such factors from CAFs which may impact tumor behavior such as the ability to metastasize, response to therapy, relapse, etc. This would aid in identifying therapeutic targets originating from the TME. Furthermore, targeting those factors along with conventional chemotherapeutic drugs is likely to enhance the overall efficacy of the therapy. This study has used fibroblasts derived from Benign Prostatic Hyperplasia (BPH) and prostate cancer for comparing the secretome using a quantitative proteomics approach. 66 proteins unique to CAFs and 24 unique to control (BPH) fibroblasts have been identified. Besides 236 proteins are differentially expressed between control and cancer-associated fibroblasts. Using in-silico approaches the potential processes that may be influenced by the differentially expressed proteins have also been identified. This study has identified both qualitative and quantitative differences between the secretomes of normal and cancer-associated fibroblasts with further validation, this paves the way for identifying therapeutic targets.

## Introduction

Epithelial tumor is not merely a mass of abnormal cells; rather a complex collection of resident and infiltrating cells, secreted factors, and the extracellular matrix (ECM). The tumor cells modify the TME to facilitate tumor growth, survival, and progression, as well as evasion of the host immune response. Tumor cells induce various molecular, cellular, and physical alterations within the tumor microenvironment (TME). TME plays a critical role in modulating tumor behavior and provides a protective niche for tumor cells. Immune cells, stromal cells, blood vessels, and ECM are typical elements of the TME (Table 1), although their composition and nature vary depending on the kind of tumor type. An interplay between tumor cells and their environment is essential to boost tumor cell heterogeneity, clonal evolution, and multidrug resistance [1, 2].

**Table 1:** Cell-types in the tumor microenvironment. This table shows the various cell-types present in the tumor microenvironment along with their major functions.

| Table 1: Cell-types in the tumor microenvironment |  |  |
| --- | --- | --- |
| Cell-type | Role | Reference |
| Regulatory T cells | Downregulate T cell induction and proliferation; contribute to poor prognosis | [32] |
| Tumor-associated Macrophages | Produce migration-stimulating factors; provide tumor cells with the ability of motility and metastasis | [33] |
| Neutrophils | Promote tumor progression and metastasis via communicating with GF's, chemokines, inflammatory factors, and other | [34] |
| Stromal cells | Helps tumor cells in immune escape and immune checkpoint blockade (ICB) resistance mechanism | [35-37] |
| Tumor Endothelial cells | Cytokines secreted by endothelial cells activate receptors on tumor cells and/or suppress anti-tumor immune reaction | [38] |
| Cancer-associated fibroblasts | ECM remodeling, angiogenesis induction, inflammatory cell recruitment, stimulate cancer cell proliferation by secre | [39,40] |
| Adipocytes | Secrete cytokines, chemokines, and hormone-like factors; obese adipose tissue hypoxia creates a highly proinflamm | [41-45] |
| Exosomes | Facilitate specific cell-cell interaction; contribute towards drug resistance; | [46,47] |
| Extracellular matrix | Facilitate seeding and metastasis of cancer cells, cause metabolic changes; acts as a barrier for drug delivery via incr | [48-50] |

As the tumor grows and progresses, the TME evolves through the secretion of various factors by cancer cells. The secreted factors i.e., the secretome can be both soluble (ex: cytokines) or insoluble factors (ex: exosomes) that are secreted into the extracellular environment [3]. The secretome is the collection of various kinds of chemokines, growth factors, coagulation factors, hormones, enzymes, cytokines, glycoproteins, and microRNAs which are secreted out of the cells. These molecules can be secreted individually or be encapsulated within extracellular vesicles or nanovesicles thereby enhancing intercellular communications [3, 4]. The secreted factors contribute to TME remodeling, fostering tumor progression by promoting angiogenesis, immune evasion, and metastatic dissemination. The Human Secretome Atlas project reports 2,641 genes encoding secreted proteins, emphasizing the complexity of secretomes irrespective of the cell types [5].

Cancer-associated fibroblasts (CAFs) are among the most prominent cell types within the TME and actively support tumor progression. These cells secrete mitogenic growth factors that promote tumor growth and influence the therapeutic response through various mechanisms [2, 5–7]. CAFs are highly heterogeneous and plastic in nature as they originate from different origins (Figure 1). CAFs and normal fibroblasts differ significantly in gene expression, phenotype, and metabolism. There are a few markers for CAFs such as alpha-smooth muscle actin (α SMA), vimentin, fibroblasts activation protein (FAP), fibroblasts specific protein 1 (FSP1), and platelet-derived growth factor receptor alpha/beta (PDGFR-α/β)[8]. However, these markers are not unique to CAFs and are usually overexpressed in CAFs [9, 10]. Despite the pleiotropic effects of CAFs on tumor behavior, very few studies have investigated their molecular distinctions. A recent study from our laboratory has identified differentially expressed mRNAs and lincRNAs between control fibroblasts vs CAFs highlighting their unique molecular profiles [11].

**Figure 1:**
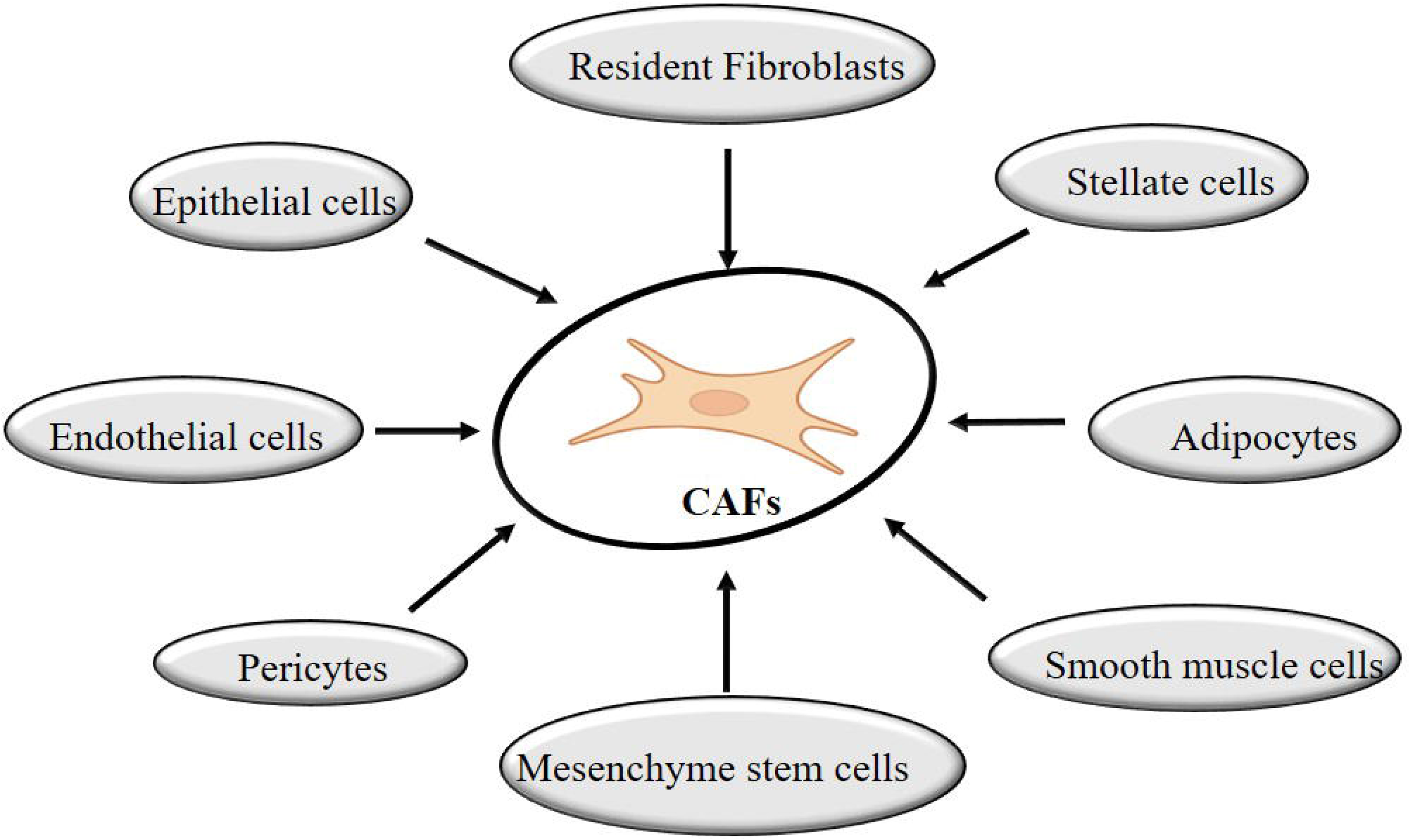
Origin of CAFs. CAFs can be derived from a variety of cell-types by trans-differentiation.

A significant amount of research and clinical trials are ongoing to understand the specific effects of TME-secreted factors on tumors, intending to use them as therapeutic targets. Fibroblast activation protein (FAP), a member of the cell surface dipeptidyl peptidase (DPP) family of serine proteases [12] is highly expressed in most cancers. An immunotoxin targeting FAP has been shown to effectively deplete FAP-positive CAFs in *in-vivo* models and exhibited potent tumor inhibitory activities across multiple cancer models [13, 14]. A multi-institutional clinical trial targeting platelet-derived growth factors (PDGF) using imatinib mesylate failed to demonstrate therapeutic benefits in patients with pancreatic ductal adenocarcinoma during phase 2 of the study (<u>NCT00161213</u>)[15]. Recombinant human hyaluronidase (PEGPH20), capable of depleting the hyaluronan in tumors, demonstrated promising results in several phase I and phase II clinical trials against pancreatic and non-small cell lung cancers [16–18]. However, in a recent phase III HALO- 301 trial, PEGPH20 failed to show efficacy against metastatic pancreatic cancer (<u>NCT02715804</u>) [19]. Yet there is limited research on the characterization of CAFs vs normal fibroblasts for their secreted factors. We conducted a label-free quantitative mass spectrometric analysis on conditioned media from control (BPH) fibroblasts and prostate cancer fibroblasts (CAFs) to address this gap. The current study aims to comprehensively identify and quantify the secreted factors, shedding light on control and CAF-specific factors that contribute to tumor cell behavior and cancer progression.

## Materials and Methods

### Clinical specimen collection

All patient samples were collected from the Institute of Nephro-Urology (Department of Urology), Bengaluru. Samples were taken either by TRUS or TURP methods. Patients who had been given a benign prostatic disease diagnosis made up the control group and were verified by a pathologist to be non-malignant. The cancer samples used in this investigation were derived from patients diagnosed with prostate cancer.

### Culturing of Fibroblasts

Fibroblasts from biopsy samples were cultured as described previously [11]. In brief, clinical specimens were transported from the hospital to the laboratory at the Centre for Human Genetics, Bengaluru in RPMI 1640 culture media supplemented with 2X PenStrep (Cat No: 15140122) and 1X GlutaMax (Cat No:350050061). The specimens were further washed with 1X PenStrep, to minimize contamination, followed by rinsing with plain RPMI 1640 (Cat No: 23400-021) growth media. Using a sterile surgical blade, the tissue samples were finely minced into small pieces as possible. These finely chopped tissue chunks were then placed into tissue culture flasks, ensuring sufficient spacing between the chunks to facilitate optimal growth.

### Secretome collection

Once cells reached 80-90% confluency, cells were washed with DPBS (Cat No: 21600-010) and incubated with serum-free media for 48 hrs. Conditioned media was collected and further processed for proteome analysis.

### Sample preparation for proteome analysis

#### Trichloroacetic acid (TCA) precipitation

Collected media was centrifuged at 500g for 5min at room temperature. The supernatant was again subjected to centrifugation at, 2000g for 10min at room temperature to remove smaller debris. Four volumes of ice-cold acetone containing 10% TCA were added to the collected supernatant and mixed by gentle vortexing. This mixture was incubated overnight at -80°C, followed by centrifugation at 13,000g for 20min at 4°C. The obtained pellet was washed with ice-cold acetone and subjected to centrifugation at 13,000g for 20min at 4°C.

The pellet obtained was dissolved in 6M guanidine hydrochloride, followed by quantification using nanodrop. 50ug of protein sample was used for further procedures. Protein samples were reduced with 5mM TCEP (tris (2-carboxyethyl) phosphine) and alkylated with 50mM iodoacetamide and then digested with trypsin (Cat No: V5280) (1:50, trypsin/lysate ratio) for 16hr at 37°C. Digests were cleaned using a C18 silica cartridge (Cat No: 89852) to remove the salt and dried using speed-vac. The dried pellet was resuspended in a buffer containing 2% acetonitrile and 0.1% formic acid. 1μg of protein was used to perform mass spectrometric analysis.

#### Mass spectrometric analysis - label-free quantification of the proteins

The mass spectrometric analysis was outsourced to BENCOS Research Solutions Pvt. Ltd, Bengaluru. After the initial quality control analysis, samples that qualified the criteria were used to carry out mass spectrometry. All the experiments were performed using an EASY-nLC 1200 system (Thermo Fisher Scientific) coupled to Thermo Fisher-QExactive plus equipped with a nanoelectrospray ion source. MS1 spectra were acquired in the Orbitrap at 70K resolution. Dynamic exclusion was employed for 10s excluding all charge states for a given precursor. MS2 spectra were acquired at 17500 resolutions (Figure 2).

**Figure 2:**
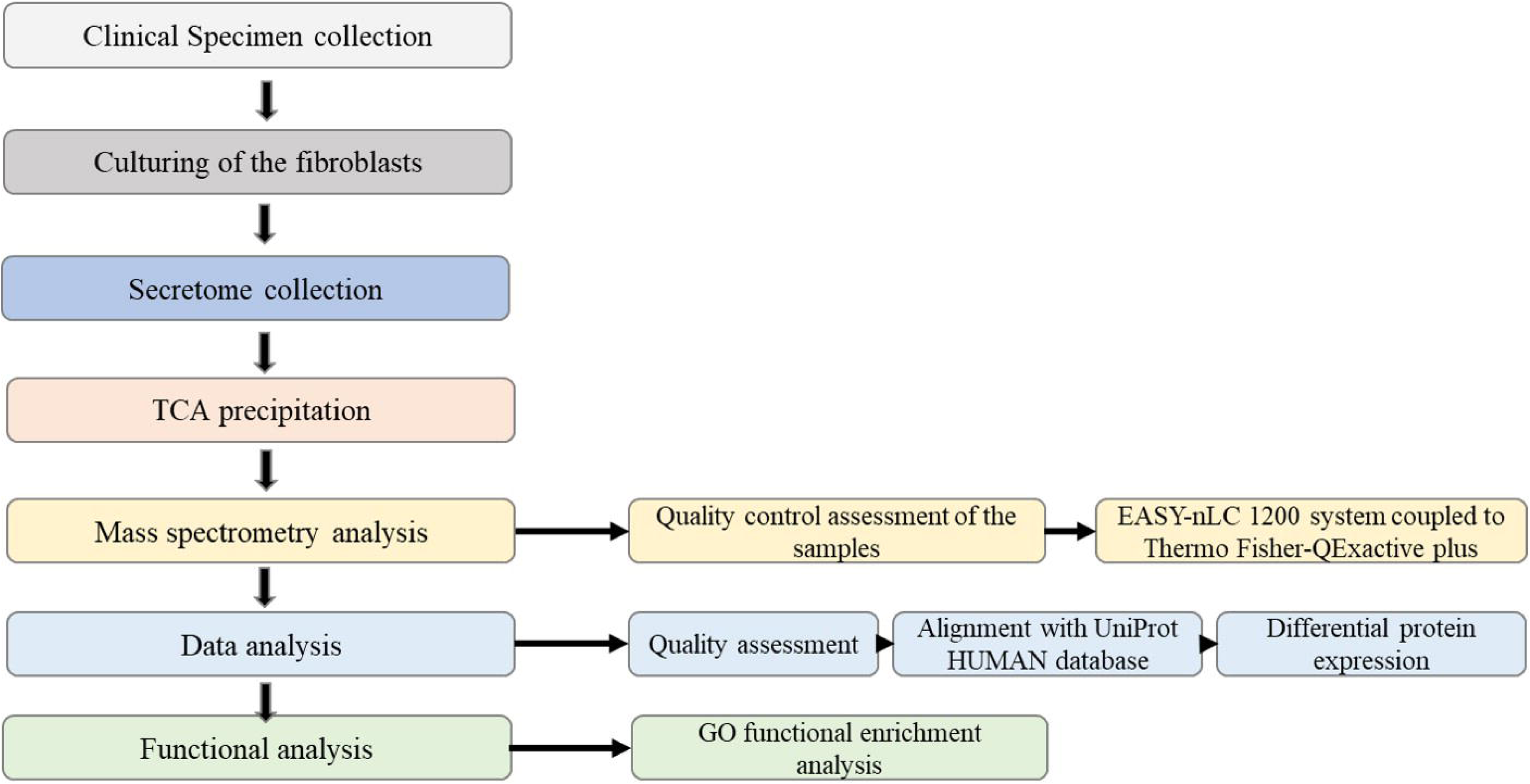
Schematic representation of the work-flow.

#### Data processing

The analysis of the label-free quantification of the proteins using Mass Spectrometry data was accomplished with the help of BENCOS Research Solutions Pvt. Ltd Bengaluru.

In brief, control samples (BPH fibroblasts) and cancer-associated fibroblasts (prostate cancer samples) were assessed for quality. High-quality reads obtained after a thorough filtering process were aligned against the UniProt HUMAN database using Thermo Proteome Discoverer (v2.4). For Sequest and Amanda’s search, the precursor and fragment mass tolerance were set at 10ppm and 0.02Da, respectively. The protease used to generate peptides, i.e. enzyme specificity was set for trypsin/P (cleavage at the C terminus of “K/R: unless followed by “P”) along with maximum missed cleavage value of two. Carbamidomethyl on cysteine as fixed modification, oxidation of methionine, and acetylation N terminus both were considered as variable modifications for database search. Both peptide spectrum match and protein false discovery rate were set to 0.01 FDR. For differential expression of proteins, log_2_>/=1.2 was used as a fold difference cut-off between CAFs and control fibroblasts.

#### Gene Ontology functional enrichment analysis

Obtained proteins were analyzed for their biological process using “enrichGO” as a function in R software (Version 4.2.0).

## Results

### Quantification of control fibroblasts and CAF proteins

A total of 1,007 proteins were identified from the secretome. The expression of these proteins varied between low to high. We have identified 236 proteins that exhibit differential expression between control fibroblasts and CAFs with a minimum log_2_ fold change of 1.2 and p-value <0.05. Of these, 119 proteins were found to be overexpressed (Table 2A), while 117 proteins were under-expressed (Table 2B) in CAFs as compared to control fibroblasts. 66 proteins unique to CAFs (Table 3A) and 24 unique to control fibroblasts (Table 3B) have been identified.

**Table 2:**
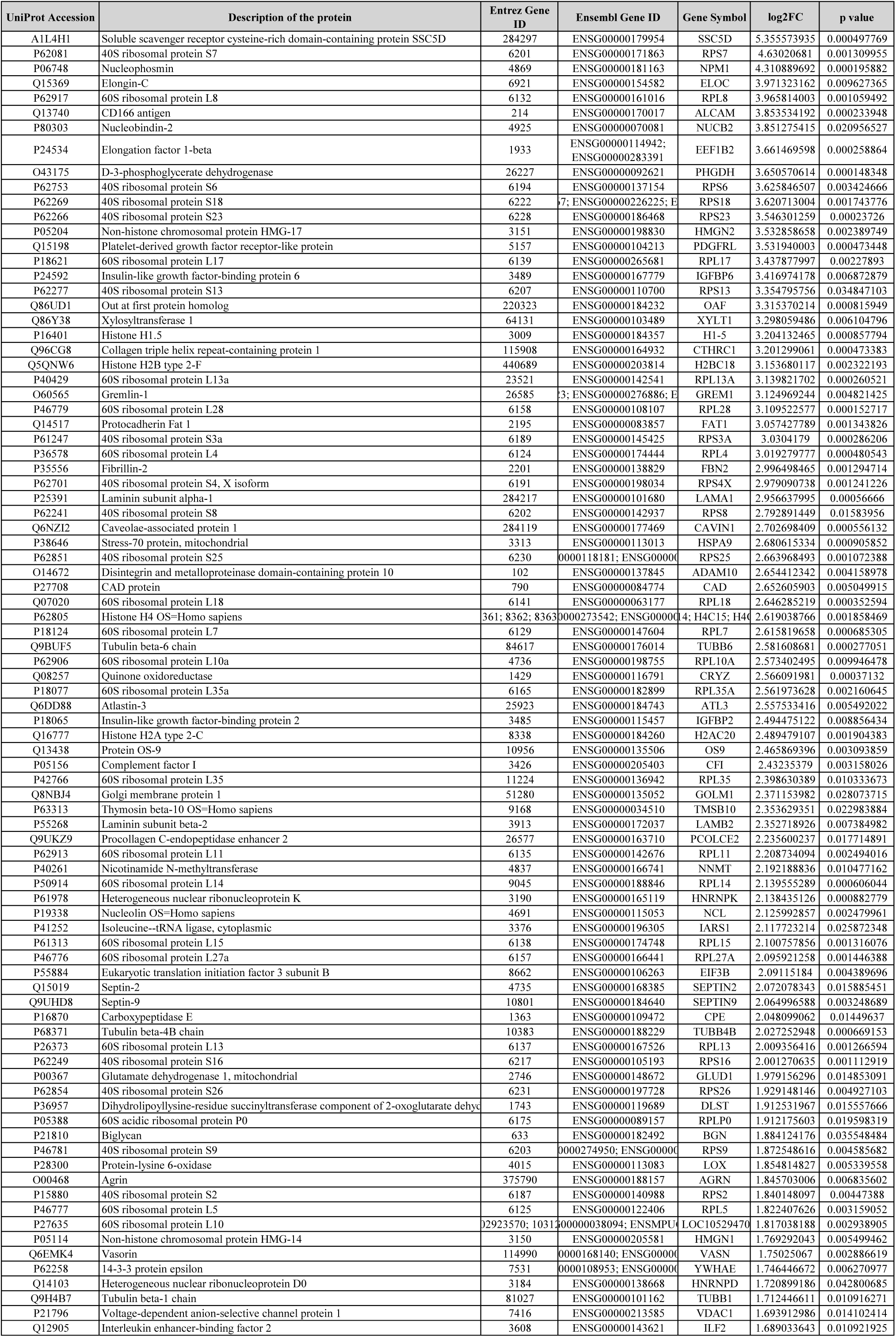
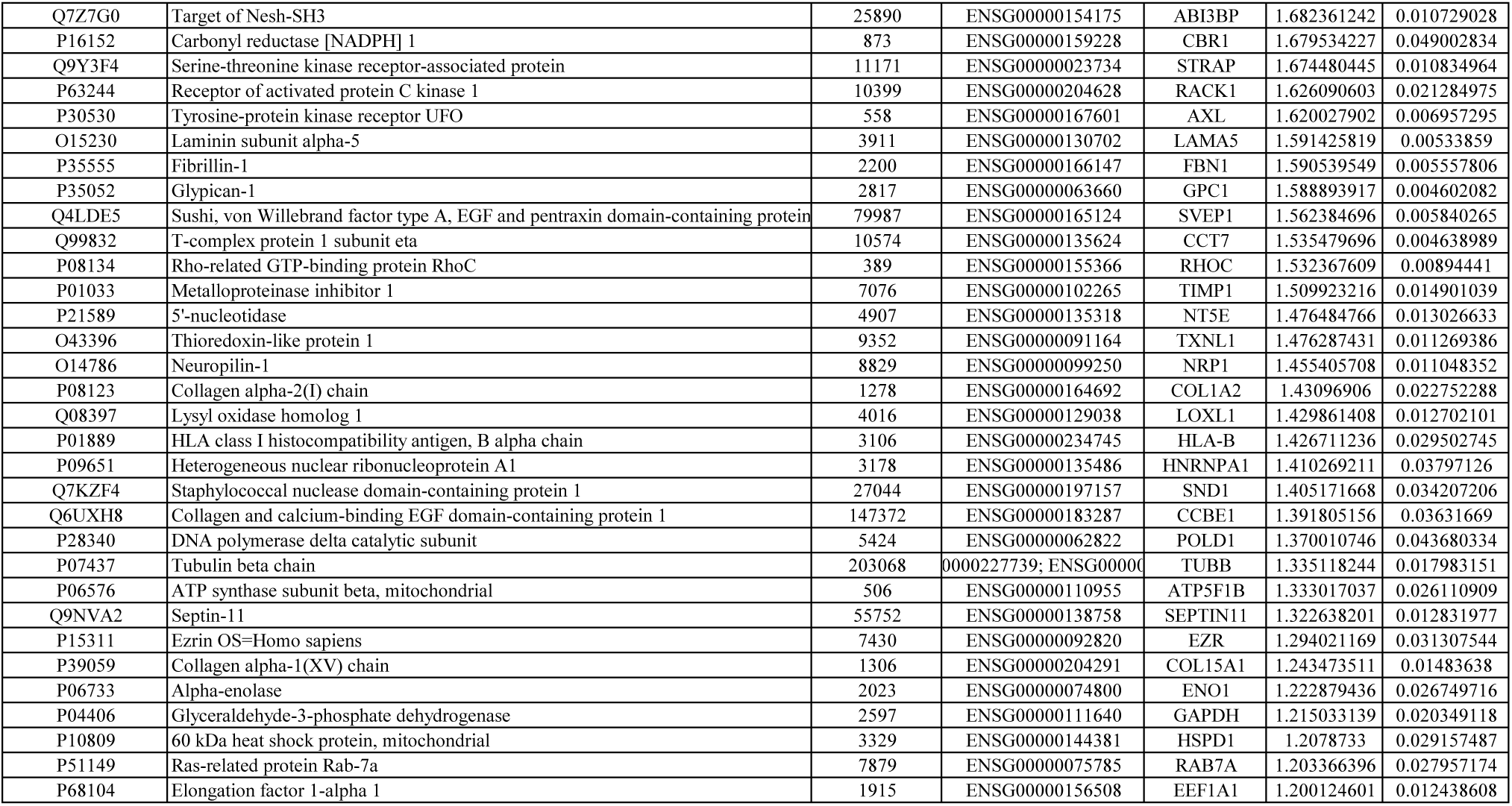
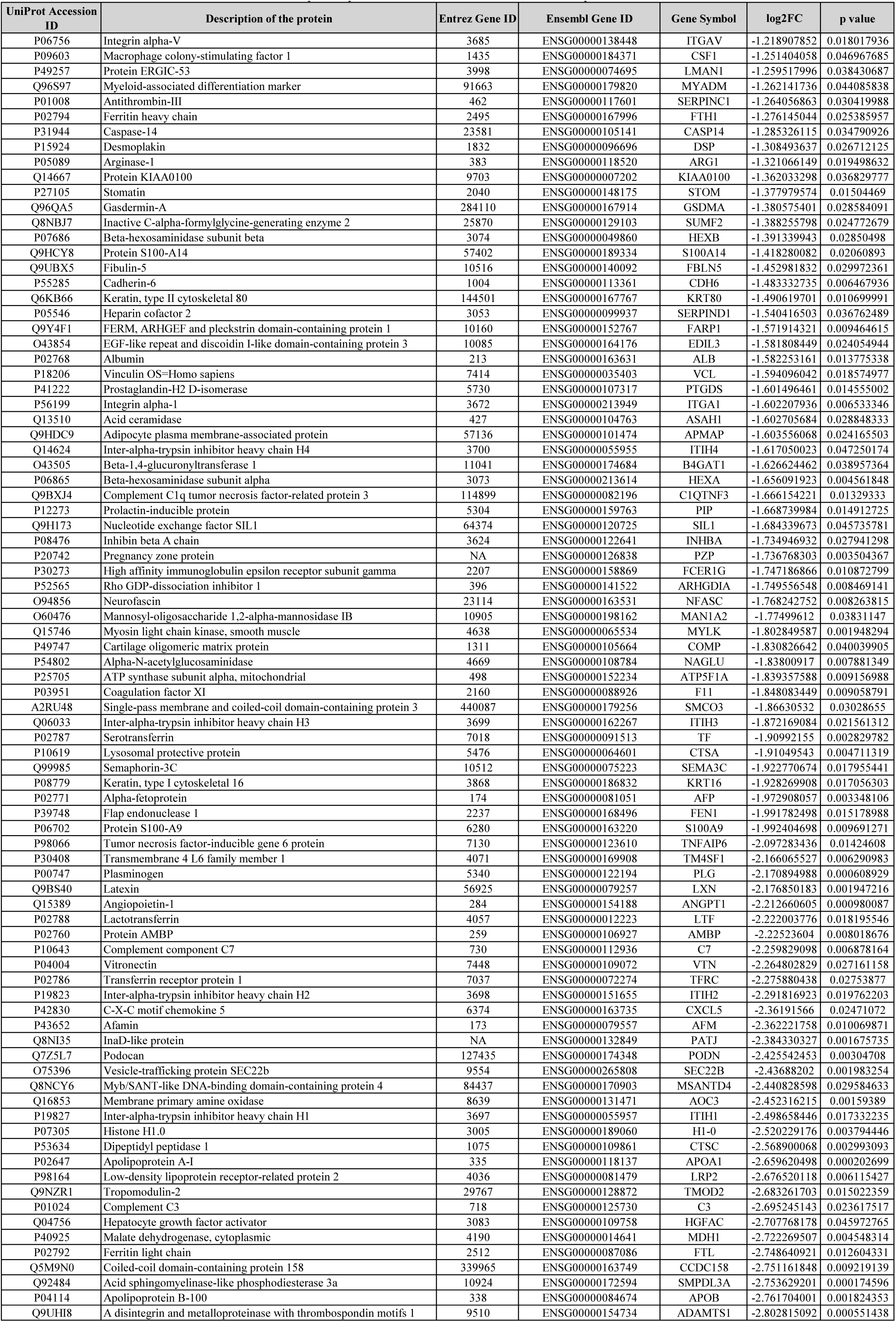
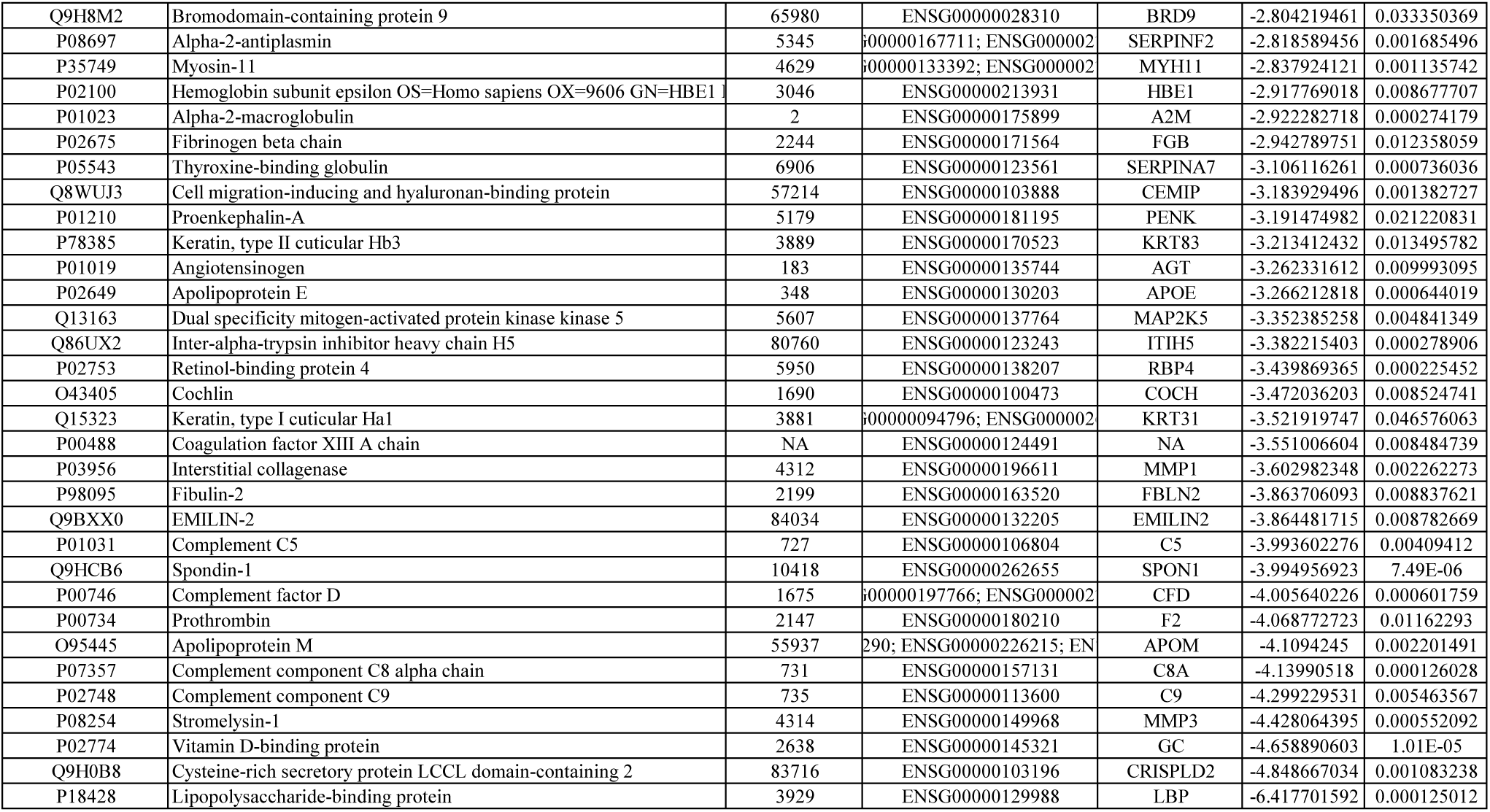
List of differentially expressed (quantitative) proteins. The list shows UniProt accession ID, description of the protein, Entrez ID, Ensembl Gene ID, gene symbol, Log_2_FC, and p-value. **A.** List of over-expressed proteins in cancer-associated fibroblasts as compared to control fibroblasts **B.** List of under-expressed proteins in cancer-associated fibroblasts as compared to control fibroblasts

**Table 3:**
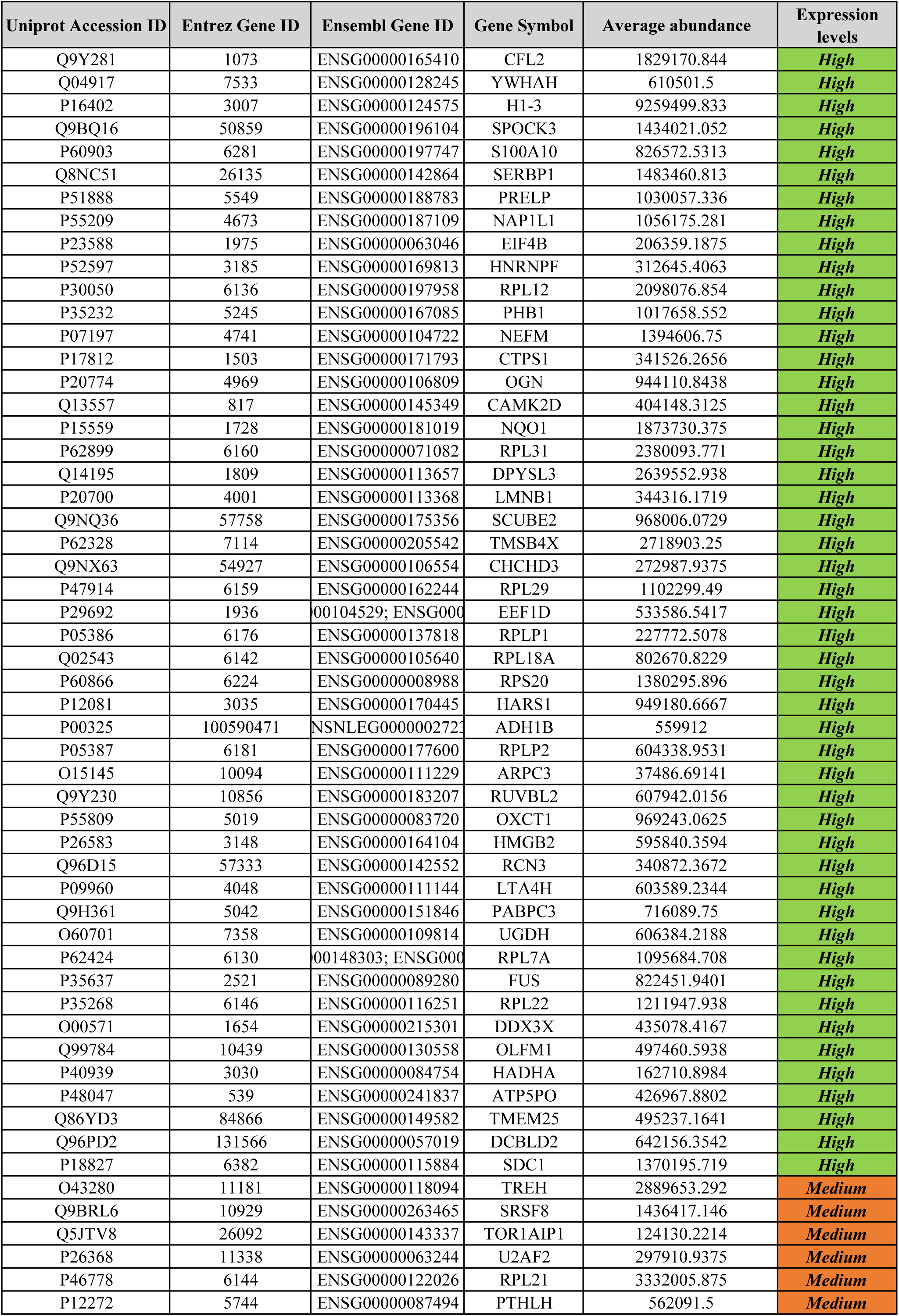
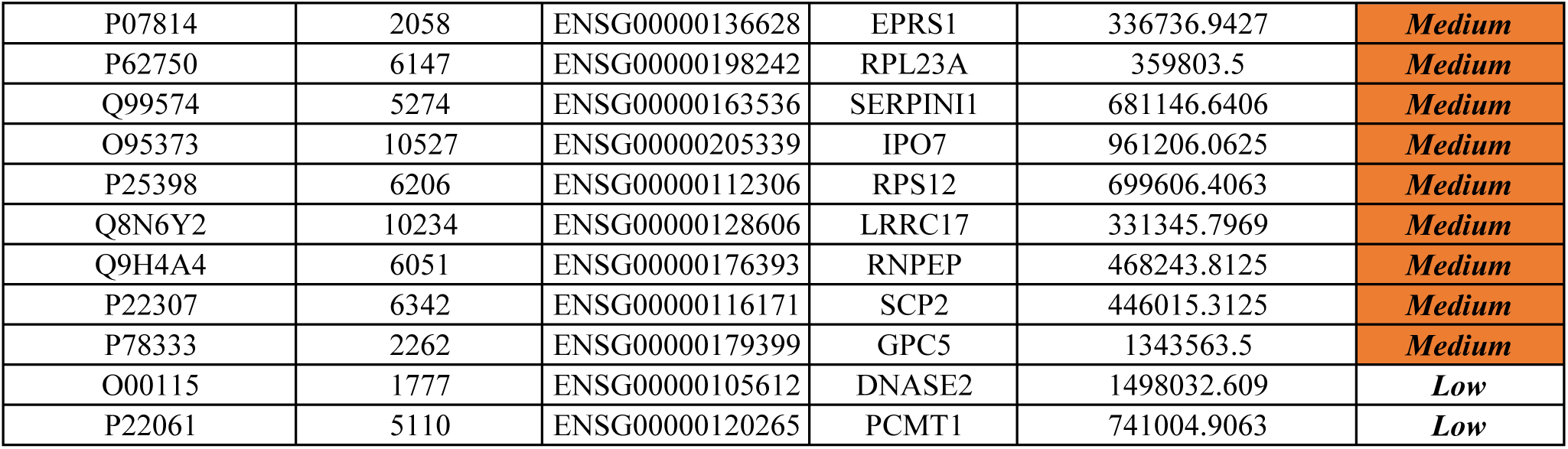
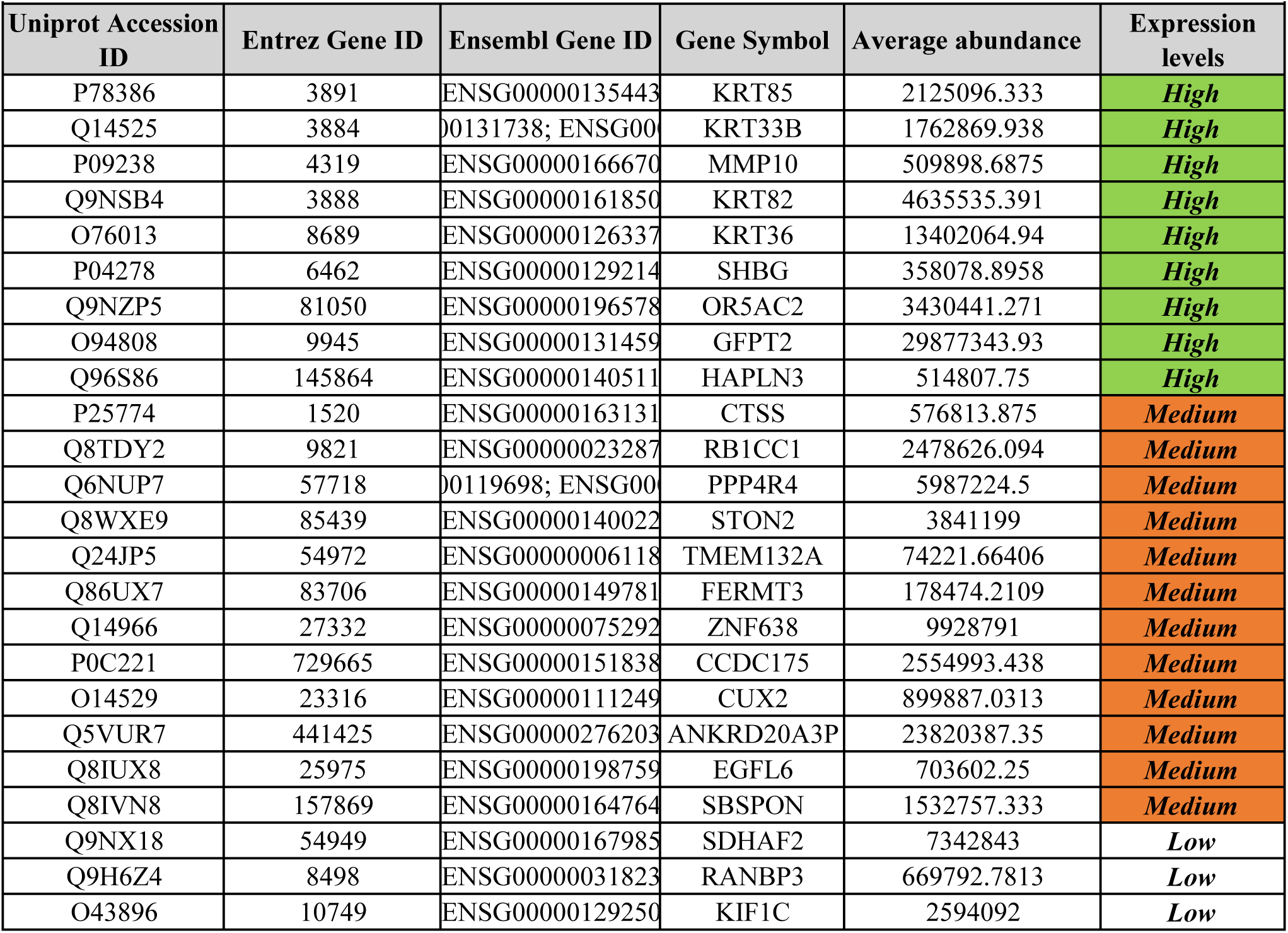
List of unique proteins expressed by cancer-associated fibroblasts and control fibroblasts. The list shows UniProt accession ID, Entrez Gene ID, Ensembl Gene ID, Gene Symbol, Average abundance, and expression levels. **A:** List of unique proteins expressed by cancer-associated fibroblasts **B:** List of unique proteins expressed by control fibroblast

### Gene Ontology functional enrichment analysis of differentially expressed proteins

Gene Ontology (GO) functional enrichment analysis for biological processes revealed that most proteins identified in the study were associated with extracellular matrix (ECM) remodeling (Figure 3) (Supplementary Table 1).

**Figure 3:**
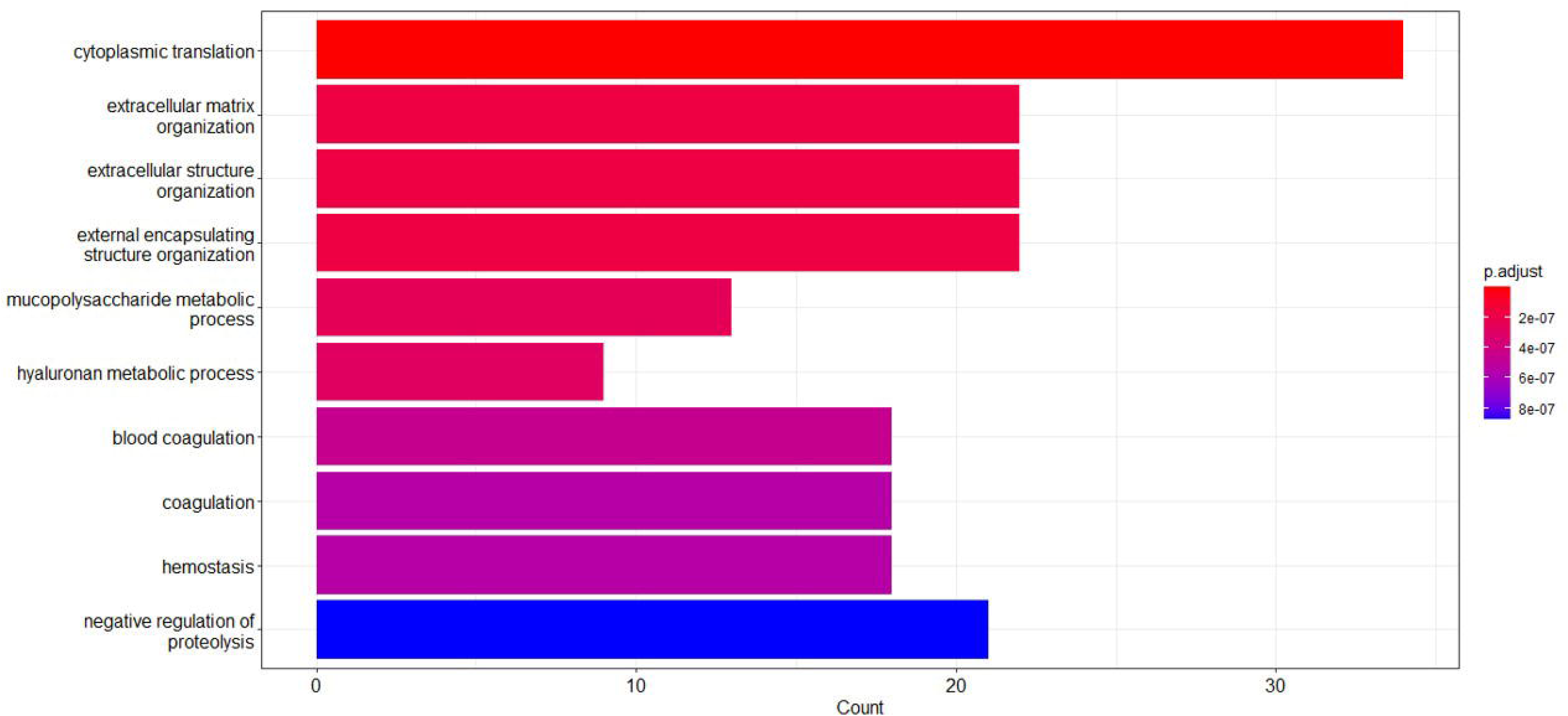
GO analysis of differentially expressed proteins.

## Discussion

The TME undergoes dynamic changes as the tumor progresses mostly due to the influence of the factors secreted by the tumor cells. The TME in turn produces factors that influence the tumor behavior [20]. CAFs are the major contributors of secreted factors. These secreted factors play a critical role in modulating tumor growth, progression, and responsiveness to therapeutic intervention.

Several studies have investigated the metabolic differences ([21–23], reviewed in [24]) and differential gene expression profiles [11, 25, 26] between normal fibroblasts and CAFs. However, relatively few studies have focused on proteomic characterization of these cells [27, 28]. Most studies available till date have identified markers which are over-expressed in CAFs. Our study compared fibroblasts obtained from malignant and non-malignant prostate as compared to most other studies which profile the CAFs in comparison to cell lines or tumors. The current study focuses on characterizing the normal and CAFs on quantitative and qualitative differences in secreted proteins. In this study, conditioned media collected from these fibroblasts were utilized for mass spectrometric analysis using label-free quantification of the proteins. This analysis has revealed several proteins unique to CAFs as well as differentially expressed between normal and cancer-associated fibroblasts. A large number of these proteins are associated with ECM remodeling. The differential quantification of these secretory factors may aid in the identification of novel biomarkers that could be explored for therapeutic targeting as well.

A comparative study conducted using conditioned media derived from CAFs obtained from metastatic colon cancer patients and normal skin fibroblasts identified an abundance of growth-promoting factors secreted by CAFs [29, 30]. Protein signatures identified through quantitative (phospho) proteomics on primary CAFs vs non-malignant prostate fibroblasts (NPF), were enriched in proteins related to cell adhesion and extracellular matrix. Additionally, the CAF proteome notably exhibited an interaction hub centered on collagen synthesis, modification, and signaling pathways. This study emphasizes the pivotal role of CAFs in ECM remodeling [28], which agrees with our observations as well. In comparison to this, our study has analyzed the secreted factors mainly because these are the key players in tumor-TME interactions.

A multiplexed mass spectrometric analysis of the secretome from colorectal cancer and lung cancer CAFs vs colorectal, lung, and pancreatic cancer cell lines, identified 2,573 secreted proteins. The study highlighted the CAF-enriched secreted proteins, particularly the members of the Wnt family and various collagen types, underscoring the potential ECM remodeling and signaling occurring within the TME [27]. Head and neck squamous carcinoma cell lines incubated with conditioned media from CAFs exhibited a robust increase in anchorage-independent growth, tumor-sphere formation, and cancer stem cell maker expression. Mass spectrometry analysis of secreted factors for normal and CAFs revealed the differential expression of epidermal growth factor receptors, insulin-like growth factor receptors, and platelet-derived growth factor receptor modulators as key paracrine cytokines. Pharmacological inhibition of these receptors resulted in a decrease in CAF secretome-induced tumorsphere formation and anchorage-independent growth. These experiments underscore the potential of CAF-secreted factors as therapeutic targets [31]. One of the upregulated genes (GREM1), and four of the downregulated genes (MMP3, PTGDS, FBLN2, APOE), identified in this study [31] agree with the findings from our current research (Supplementary Table 2). This study utilized a single population of CAFs and normal fibroblasts. In contrast, our study has used three distinct populations of CAFs and control fibroblasts, providing a more comprehensive and representative analysis. Taken together our findings provide a foundation for understanding the differential expression of the secreted proteins between control fibroblasts and CAFs. These insights could serve as a basis for identifying therapeutic targets, ultimately aiding in overcoming TME-mediated tumor progression and resistance to therapy.

## Authors contribution

AA analyzed the proteomics data and prepared the manuscript draft with all figures and tables. MSM, RRA, and NN helped in the collection of patient samples, deriving the fibroblasts, and preliminary characterization.

VB and NT helped in the collection of patient samples and clinical/pathological evaluation. RK helped co-ordinate all the patient-related work and helped in procuring funding.

PR conceived and strategized the study, procured the funding, and finalized the manuscript. All authors reviewed and approved the manuscript

## Ethics approval and consent to participate

The study was approved by the Institutional Ethics Committee of both the participating institutions (CHG/077(b)/IEC/2019-20/001 and EC/01/2019). Informed consent has been obtained from all participants whose tissue samples have been used in this study. The identity of the patients has been kept confidential.

The study has been conducted in accordance with the Declaration of Helsinki.

This study presented here was funded by the Indian Council for Medical Research, Govt of India (2019 − 0937), granted to PR.

## Supplementary Table legends

**Supplementary Table 1:** List of secreted proteins and the biological processes they are involved in

The list shows GO ID, description, p-value, p-adjust, q-value, Gene ID, and count.

**Supplementary Table 2:** Comparison of data with published data

The shows UniProt accession ID, Description of the protein, Entrez Gene ID, Ensembl Gene ID, Gene Symbol, log_2_FC from the current study, p-value from the current study, log_2_FC from the available study, p-value from the available study.

