## Supplementary tables for "Secretome analysis of cancer-associated fibroblasts from prostate cancer to identify potential therapeutic targets"

| Supplementary Table 1: List of secreted proteins and the biological processes they are involved in | | | | | | |
|---|---|---|---|---|---|---|
| **GO ID** | **Description** | **pvalue** | **p.adjust** | **qvalue** | **GeneID** | **Count** |
| GO:0002181 | cytoplasmic translation | 1.06E-31 | 3.65E-28 | 3.06E-28 | RPLP0/RPS2/RPL7/RP | 34 |
| GO:0030198 | extracellular matrix organization | 1.68E-10 | 1.74E-07 | 1.46E-07 | ADAM10/GREM1/AG | 22 |
| GO:0043062 | extracellular structure organization | 1.79E-10 | 1.74E-07 | 1.46E-07 | ADAM10/GREM1/AG | 22 |
| GO:0045229 | external encapsulating structure organization | 2.02E-10 | 1.74E-07 | 1.46E-07 | ADAM10/GREM1/AG | 22 |
| GO:1903510 | mucopolysaccharide metabolic process | 3.59E-10 | 2.48E-07 | 2.08E-07 | B4GAT1/HEXA/HEXE | 13 |
| GO:0030212 | hyaluronan metabolic process | 4.99E-10 | 2.88E-07 | 2.41E-07 | HEXA/HEXB/ITIH2/IT | 9 |
| GO:0007596 | blood coagulation | 9.28E-10 | 4.59E-07 | 3.84E-07 | F2/SERPINC1/APOE/F | 18 |
| GO:0050817 | coagulation | 1.32E-09 | 5.46E-07 | 4.57E-07 | F2/SERPINC1/APOE/F | 18 |
| GO:0007599 | hemostasis | 1.42E-09 | 5.46E-07 | 4.57E-07 | F2/SERPINC1/APOE/F | 18 |
| GO:0045861 | negative regulation of proteolysis | 2.53E-09 | 8.77E-07 | 7.34E-07 | F2/A2M/TIMP1/LTF/V | 21 |
| GO:0030203 | glycosaminoglycan metabolic process | 4.32E-09 | 1.36E-06 | 1.14E-06 | B4GAT1/HEXA/HEXE | 13 |
| GO:0006022 | aminoglycan metabolic process | 1.07E-08 | 3.10E-06 | 2.59E-06 | B4GAT1/HEXA/HEXE | 13 |
| GO:0010466 | negative regulation of peptidase activity | 1.43E-08 | 3.81E-06 | 3.19E-06 | A2M/TIMP1/LTF/VTN | 17 |
| GO:0042060 | wound healing | 1.73E-08 | 4.28E-06 | 3.58E-06 | F2/SERPINC1/TIMP1/ | 23 |
| GO:1900046 | regulation of hemostasis | 3.15E-08 | 7.25E-06 | 6.07E-06 | F2/SERPINC1/APOE/F | 10 |
| GO:0006959 | humoral immune response | 3.35E-08 | 7.25E-06 | 6.07E-06 | F2/CFD/A2M/C3/C5/F | 17 |
| GO:0031589 | cell-substrate adhesion | 5.69E-08 | 1.11E-05 | 9.33E-06 | NRP1/LAMA5/EDIL3/ | 20 |
| GO:0006956 | complement activation | 5.80E-08 | 1.11E-05 | 9.33E-06 | CFD/A2M/C3/C5/C9/C | 9 |
| GO:0006957 | complement activation, alternative pathway | 7.58E-08 | 1.38E-05 | 1.16E-05 | CFD/C3/C5/C9/C8A/C | 6 |
| GO:0051235 | maintenance of location | 1.62E-07 | 2.80E-05 | 2.34E-05 | F2/C3/APOA1/APOE/ | 19 |
| GO:0006958 | complement activation, classical pathway | 1.90E-07 | 3.12E-05 | 2.62E-05 | C3/C5/C9/CFI/C8A/C7/ | 7 |
| GO:1900047 | negative regulation of hemostasis | 2.95E-07 | 4.64E-05 | 3.88E-05 | F2/APOE/FGB/VTN/SI | 8 |
| GO:0030193 | regulation of blood coagulation | 3.09E-07 | 4.65E-05 | 3.89E-05 | F2/SERPINC1/APOE/F | 9 |
| GO:0050818 | regulation of coagulation | 5.64E-07 | 8.13E-05 | 6.80E-05 | F2/SERPINC1/APOE/F | 9 |
| GO:0051346 | negative regulation of hydrolase activity | 6.99E-07 | 9.68E-05 | 8.10E-05 | A2M/TIMP1/APOA1/L | 18 |
| GO:0052547 | regulation of peptidase activity | 8.44E-07 | 0.000112 | 9.40E-05 | A2M/TIMP1/LTF/VTN | 20 |
| GO:0042730 | fibrinolysis | 1.09E-06 | 0.00014 | 0.000117 | F2/FGB/VTN/SERPINI | 6 |
| GO:0050878 | regulation of body fluid levels | 1.34E-06 | 0.000166 | 0.000139 | F2/SERPINC1/APOE/F | 18 |
| GO:0002455 | humoral immune response mediated by circulating immun | 2.06E-06 | 0.000243 | 0.000204 | C3/C5/C9/CFI/C8A/C7/ | 7 |

| | | | | | | |
|---|---|---|---|---|---|---|
| GO:0007229 | integrin-mediated signaling pathway | 2.11E-06 | 0.000243 | 0.000204 | ADAM10/NRP1/LAM/ | 10 |
| GO:0007568 | aging | 2.33E-06 | 0.00026 | 0.000218 | AGT/TIMP1/PENK/TF | 12 |
| GO:0051917 | regulation of fibrinolysis | 2.71E-06 | 0.000294 | 0.000246 | F2/VTN/SERPINF2/PL | 5 |
| GO:0042273 | ribosomal large subunit biogenesis | 3.16E-06 | 0.000332 | 0.000278 | RPLP0/NPM1/RPL7/RI | 8 |
| GO:0030195 | negative regulation of blood coagulation | 3.67E-06 | 0.000374 | 0.000313 | F2/APOE/FGB/VTN/SI | 7 |
| GO:0010811 | positive regulation of cell-substrate adhesion | 5.70E-06 | 0.000549 | 0.000459 | NRP1/EDIL3/APOA1/I | 10 |
| GO:0045785 | positive regulation of cell adhesion | 5.71E-06 | 0.000549 | 0.000459 | NRP1/EDIL3/APOA1/I | 20 |
| GO:0050819 | negative regulation of coagulation | 6.22E-06 | 0.000569 | 0.000476 | F2/APOE/FGB/VTN/SI | 7 |
| GO:0060445 | branching involved in salivary gland morphogenesis | 6.24E-06 | 0.000569 | 0.000476 | NRP1/LAMA5/TGM2/ | 5 |
| GO:0032102 | negative regulation of response to external stimulus | 6.78E-06 | 0.000602 | 0.000504 | NRP1/GREM1/F2/A2M | 19 |
| GO:0010810 | regulation of cell-substrate adhesion | 7.44E-06 | 0.000643 | 0.000538 | NRP1/EDIL3/GREM1/ | 13 |
| GO:0030168 | platelet activation | 9.90E-06 | 0.00083 | 0.000694 | F2/APOE/FGB/VCL/A | 10 |
| GO:0002449 | lymphocyte mediated immunity | 1.01E-05 | 0.00083 | 0.000694 | C3/C5/HLA-B/C9/TFR | 15 |
| GO:0061041 | regulation of wound healing | 1.29E-05 | 0.001035 | 0.000866 | F2/SERPINC1/APOE/F | 10 |
| GO:0034113 | heterotypic cell-cell adhesion | 1.42E-05 | 0.001116 | 0.000934 | APOA1/FGB/ITGAV/J | 7 |
| GO:0060326 | cell chemotaxis | 1.60E-05 | 0.001231 | 0.00103 | ADAM10/NRP1/GREM | 15 |
| GO:0044403 | biological process involved in symbiotic interaction | 1.66E-05 | 0.00125 | 0.001047 | NRP1/F2/APOE/TFRC/ | 15 |
| GO:0030595 | leukocyte chemotaxis | 1.79E-05 | 0.001317 | 0.001103 | ADAM10/GREM1/C5/ | 13 |
| GO:0051918 | negative regulation of fibrinolysis | 1.85E-05 | 0.001332 | 0.001115 | F2/VTN/SERPINF2/PL | 4 |
| GO:0016064 | immunoglobulin mediated immune response | 2.11E-05 | 0.001491 | 0.001248 | C3/C5/C9/TFRC/CFI/C | 10 |
| GO:0002443 | leukocyte mediated immunity | 2.22E-05 | 0.00154 | 0.001289 | F2/C3/C5/HLA-B/C9/T | 17 |
| GO:0051702 | biological process involved in interaction with symbiont | 2.47E-05 | 0.001675 | 0.001402 | F2/APOE/LTF/GAPDH | 9 |
| GO:0019724 | B cell mediated immunity | 2.52E-05 | 0.001677 | 0.001404 | C3/C5/C9/TFRC/CFI/C | 10 |
| GO:0048771 | tissue remodeling | 2.68E-05 | 0.001748 | 0.001463 | GREM1/AGT/TIMP1/T | 11 |
| GO:0030194 | positive regulation of blood coagulation | 2.80E-05 | 0.001761 | 0.001474 | F2/VTN/SERPINF2/EM | 5 |
| GO:1900048 | positive regulation of hemostasis | 2.80E-05 | 0.001761 | 0.001474 | F2/VTN/SERPINF2/EM | 5 |
| GO:0006898 | receptor-mediated endocytosis | 3.37E-05 | 0.002081 | 0.001741 | GREM1/SNAP91/C3/A | 13 |
| GO:0016072 | rRNA metabolic process | 3.51E-05 | 0.002128 | 0.001781 | RPL7/NCL/RPL35/RPL | 13 |
| GO:0006953 | acute-phase response | 3.61E-05 | 0.002154 | 0.001803 | F2/A2M/TFRC/SERPIN | 6 |
| GO:0045807 | positive regulation of endocytosis | 3.71E-05 | 0.002161 | 0.001809 | GREM1/C3/APOE/TF/ | 8 |
| GO:0042254 | ribosome biogenesis | 3.96E-05 | 0.002161 | 0.001809 | RPLP0/NPM1/RPL7/RI | 14 |
| GO:0034368 | protein-lipid complex remodeling | 3.97E-05 | 0.002161 | 0.001809 | APOM/AGT/APOA1/A | 5 |

| | | | | | | |
|---|---|---|---|---|---|---|
| GO:0034369 | plasma lipoprotein particle remodeling | 3.97E-05 | 0.002161 | 0.001809 | APOM/AGT/APOA1/A | 5 |
| GO:0050820 | positive regulation of coagulation | 3.97E-05 | 0.002161 | 0.001809 | F2/VTN/SERPINF2/EM | 5 |
| GO:0034109 | homotypic cell-cell adhesion | 4.00E-05 | 0.002161 | 0.001809 | FGB/JUP/DSP/VCL/C( | 8 |
| GO:0045109 | intermediate filament organization | 4.64E-05 | 0.002471 | 0.002068 | KRT16/KRT5/DSP/KR | 7 |
| GO:0007435 | salivary gland morphogenesis | 5.49E-05 | 0.002838 | 0.002375 | NRP1/LAMA5/TGM2/ | 5 |
| GO:0034367 | protein-containing complex remodeling | 5.49E-05 | 0.002838 | 0.002375 | APOM/AGT/APOA1/A | 5 |
| GO:0071621 | granulocyte chemotaxis | 5.64E-05 | 0.002838 | 0.002376 | CSF1/LBP/ITGA1/PPI/ | 9 |
| GO:0043588 | skin development | 5.66E-05 | 0.002838 | 0.002376 | LAMA5/COL1A2/KRT | 14 |
| GO:0001906 | cell killing | 5.81E-05 | 0.002873 | 0.002405 | F2/C3/HLA-B/C9/LTF/( | 11 |
| GO:0051651 | maintenance of location in cell | 5.90E-05 | 0.002875 | 0.002406 | F2/APOE/ALB/FTL/FT | 12 |
| GO:0048260 | positive regulation of receptor-mediated endocytosis | 6.39E-05 | 0.003074 | 0.002573 | GREM1/C3/TF/VTN/H | 6 |
| GO:0042274 | ribosomal small subunit biogenesis | 7.12E-05 | 0.00333 | 0.002787 | NPM1/RPS9/RPS7/RPS | 7 |
| GO:0061045 | negative regulation of wound healing | 7.12E-05 | 0.00333 | 0.002787 | F2/APOE/FGB/VTN/SI | 7 |
| GO:0007431 | salivary gland development | 7.43E-05 | 0.003429 | 0.00287 | NRP1/LAMA5/TGM2/ | 5 |
| GO:0097529 | myeloid leukocyte migration | 8.15E-05 | 0.003676 | 0.003077 | GREM1/C5/CSF1/LBP/ | 12 |
| GO:1903034 | regulation of response to wounding | 8.18E-05 | 0.003676 | 0.003077 | F2/SERPINC1/APOE/F | 10 |
| GO:0060840 | artery development | 9.19E-05 | 0.004077 | 0.003412 | NRP1/APOE/APOB/L( | 8 |
| GO:0016049 | cell growth | 9.40E-05 | 0.004118 | 0.003446 | ADAM10/NRP1/GREM | 18 |
| GO:0001649 | osteoblast differentiation | 0.000115 | 0.004988 | 0.004175 | GREM1/PENK/LTF/AT | 12 |
| GO:0019915 | lipid storage | 0.000133 | 0.005647 | 0.004726 | C3/APOA1/APOE/APC | 7 |
| GO:0048259 | regulation of receptor-mediated endocytosis | 0.000134 | 0.005647 | 0.004726 | GREM1/SNAP91/C3/T | 8 |
| GO:0030728 | ovulation | 0.000142 | 0.005905 | 0.004943 | AFP/TNFAIP6/ADAM | 4 |
| GO:0006364 | rRNA processing | 0.000143 | 0.005905 | 0.004943 | RPL7/RPL35/RPL14/RI | 11 |
| GO:0090303 | positive regulation of wound healing | 0.000155 | 0.006322 | 0.005292 | F2/VTN/SERPINF2/M\ | 6 |
| GO:0001503 | ossification | 0.000173 | 0.006971 | 0.005834 | GREM1/PENK/LTF/AT | 16 |
| GO:0022613 | ribonucleoprotein complex biogenesis | 0.000178 | 0.007071 | 0.005918 | RPLP0/NPM1/RPL7/RI | 17 |
| GO:0001894 | tissue homeostasis | 0.000185 | 0.007126 | 0.005964 | RBP4/ALB/TFRC/TF/L | 12 |
| GO:0060249 | anatomical structure homeostasis | 0.000185 | 0.007126 | 0.005964 | RBP4/ALB/TFRC/TF/L | 12 |
| GO:0030199 | collagen fibril organization | 0.000185 | 0.007126 | 0.005964 | GREM1/COL1A2/SER | 6 |
| GO:0046660 | female sex differentiation | 0.000201 | 0.007611 | 0.00637 | A2M/RBP4/AFP/AXL/ | 8 |
| GO:0045104 | intermediate filament cytoskeleton organization | 0.000202 | 0.007611 | 0.00637 | KRT16/KRT5/DSP/KR | 7 |
| GO:0019835 | cytolysis | 0.000206 | 0.007642 | 0.006396 | C5/C9/C8A/C7 | 4 |

| | | | | | | |
|---|---|---|---|---|---|---|
| GO:0085029 | extracellular matrix assembly | 0.000209 | 0.007642 | 0.006396 | AGT/COL1A2/LOX/M | 5 |
| GO:0022612 | gland morphogenesis | 0.000213 | 0.007642 | 0.006396 | NRP1/LAMA5/CSF1/T | 8 |
| GO:0097530 | granulocyte migration | 0.000214 | 0.007642 | 0.006396 | CSF1/LBP/ITGA1/PPI | 9 |
| GO:0045103 | intermediate filament-based process | 0.000216 | 0.007642 | 0.006396 | KRT16/KRT5/DSP/KR | 7 |
| GO:1903035 | negative regulation of response to wounding | 0.000216 | 0.007642 | 0.006396 | F2/APOE/FGB/VTN/S | 7 |
| GO:0002460 | adaptive immune response based on somatic recombinatio | 0.000238 | 0.00831 | 0.006955 | C3/C5/HLA-B/C9/TFR | 13 |
| GO:0048251 | elastic fiber assembly | 0.000246 | 0.008361 | 0.006998 | LOX/MYH11/FBLN5 | 3 |
| GO:0060068 | vagina development | 0.000246 | 0.008361 | 0.006998 | RBP4/AXL/LRP2 | 3 |
| GO:1905918 | regulation of CoA-transferase activity | 0.000246 | 0.008361 | 0.006998 | AGT/APOA1/APOE | 3 |
| GO:0019730 | antimicrobial humoral response | 0.000251 | 0.008444 | 0.007067 | F2/FGB/TF/LTF/GAPD | 8 |
| GO:0010951 | negative regulation of endopeptidase activity | 0.000259 | 0.00862 | 0.007215 | A2M/TIMP1/LTF/VTN | 9 |
| GO:0034114 | regulation of heterotypic cell-cell adhesion | 0.000289 | 0.009522 | 0.00797 | APOA1/FGB/MYADM | 4 |
| GO:0007160 | cell-matrix adhesion | 0.000318 | 0.009928 | 0.00831 | NRP1/GREM1/FGB/V | 11 |
| GO:0014911 | positive regulation of smooth muscle cell migration | 0.000322 | 0.009928 | 0.00831 | NRP1/AGT/VTN/ADA | 5 |
| GO:0035272 | exocrine system development | 0.000322 | 0.009928 | 0.00831 | NRP1/LAMA5/TGM2/ | 5 |
| GO:0071827 | plasma lipoprotein particle organization | 0.000322 | 0.009928 | 0.00831 | APOM/AGT/APOA1/A | 5 |
| GO:0034115 | negative regulation of heterotypic cell-cell adhesion | 0.000336 | 0.009928 | 0.00831 | APOA1/MYADM/MA | 3 |
| GO:0035581 | sequestering of extracellular ligand from receptor | 0.000336 | 0.009928 | 0.00831 | GREM1/FBN1/FBN2 | 3 |
| GO:0070587 | regulation of cell-cell adhesion involved in gastrulation | 0.000336 | 0.009928 | 0.00831 | APOA1/MYADM/MA | 3 |
| GO:0071492 | cellular response to UV-A | 0.000336 | 0.009928 | 0.00831 | TIMP1/MMP1/MMP3 | 3 |
| GO:0031639 | plasminogen activation | 0.000338 | 0.009928 | 0.00831 | FGB/ENO1/SERPINF2 | 4 |
| GO:0034377 | plasma lipoprotein particle assembly | 0.000338 | 0.009928 | 0.00831 | APOM/APOA1/APOE | 4 |
| GO:0051238 | sequestering of metal ion | 0.000338 | 0.009928 | 0.00831 | FTL/FTH1/LMAN1/S1 | 4 |
| GO:0060343 | trabecula formation | 0.000338 | 0.009928 | 0.00831 | GREM1/RBP4/FBN2/A | 4 |
| GO:0034446 | substrate adhesion-dependent cell spreading | 0.000338 | 0.009928 | 0.00831 | NRP1/LAMA5/APOA1 | 7 |
| GO:0043542 | endothelial cell migration | 0.000361 | 0.010467 | 0.008761 | NRP1/GREM1/AGT/A | 12 |
| GO:1904951 | positive regulation of establishment of protein localization | 0.000363 | 0.010467 | 0.008761 | GLUD1/F2/FGB/RBP4 | 13 |
| GO:0070527 | platelet aggregation | 0.000382 | 0.010916 | 0.009136 | FGB/VCL/COMP/PPIA | 6 |
| GO:0071825 | protein-lipid complex subunit organization | 0.000434 | 0.012172 | 0.010188 | APOM/AGT/APOA1/A | 5 |
| GO:0001558 | regulation of cell growth | 0.000435 | 0.012172 | 0.010188 | ADAM10/NRP1/GREM | 15 |
| GO:0042698 | ovulation cycle | 0.000442 | 0.012172 | 0.010188 | A2M/AFP/AXL/TNFA | 6 |
| GO:0070586 | cell-cell adhesion involved in gastrulation | 0.000443 | 0.012172 | 0.010188 | APOA1/MYADM/MA | 3 |

| | | | | | | |
|---|---|---|---|---|---|---|
| GO:1904667 | negative regulation of ubiquitin protein ligase activity | 0.000443 | 0.012172 | 0.010188 | RPS7/RPL11/RPL5 | 3 |
| GO:1903036 | positive regulation of response to wounding | 0.000474 | 0.012929 | 0.010822 | F2/VTN/SERPINF2/M | 6 |
| GO:0050821 | protein stabilization | 0.000507 | 0.013592 | 0.011376 | APOA1/GAPDH/HSPI | 10 |
| GO:2001233 | regulation of apoptotic signaling pathway | 0.000509 | 0.013592 | 0.011376 | NRP1/AGT/FGB/ENO | 14 |
| GO:0030593 | neutrophil chemotaxis | 0.000511 | 0.013592 | 0.011376 | LBP/ITGA1/PPIA/TNF | 7 |
| GO:0065005 | protein-lipid complex assembly | 0.000521 | 0.013755 | 0.011513 | APOM/APOA1/APOE | 4 |
| GO:0030100 | regulation of endocytosis | 0.000526 | 0.013779 | 0.011533 | GREM1/SNAP91/C3/A | 10 |
| GO:0032963 | collagen metabolic process | 0.00054 | 0.014051 | 0.01176 | F2/MMP1/COL1A2/MI | 7 |
| GO:0033690 | positive regulation of osteoblast proliferation | 0.000571 | 0.014384 | 0.012039 | LTF/ITGAV/CTHRC1 | 3 |
| GO:0034380 | high-density lipoprotein particle assembly | 0.000571 | 0.014384 | 0.012039 | APOM/APOA1/APOE | 3 |
| GO:1900115 | extracellular regulation of signal transduction | 0.000571 | 0.014384 | 0.012039 | GREM1/FBN1/FBN2 | 3 |
| GO:1900116 | extracellular negative regulation of signal transduction | 0.000571 | 0.014384 | 0.012039 | GREM1/FBN1/FBN2 | 3 |
| GO:0022602 | ovulation cycle process | 0.000574 | 0.014384 | 0.012039 | A2M/AFP/TNFAIP6/A | 5 |
| GO:0043534 | blood vessel endothelial cell migration | 0.000589 | 0.014664 | 0.012273 | NRP1/GREM1/APOA1 | 9 |
| GO:0048844 | artery morphogenesis | 0.000625 | 0.015439 | 0.012922 | NRP1/APOE/APOB/C | 6 |
| GO:0090287 | regulation of cellular response to growth factor stimulus | 0.000636 | 0.015622 | 0.013076 | NRP1/GREM1/AGT/V | 13 |
| GO:0033688 | regulation of osteoblast proliferation | 0.000676 | 0.01647 | 0.013786 | GREM1/LTF/ITGAV/C | 4 |
| GO:0002683 | negative regulation of immune system process | 0.000698 | 0.016905 | 0.01415 | GREM1/A2M/C5/HLA | 16 |
| GO:0061844 | antimicrobial humoral immune response mediated by antin | 0.000712 | 0.017047 | 0.014268 | F2/LTF/GAPDH/HMG | 6 |
| GO:0018158 | protein oxidation | 0.000719 | 0.017047 | 0.014268 | APOA1/LOX/LOXL1 | 3 |
| GO:0070141 | response to UV-A | 0.000719 | 0.017047 | 0.014268 | TIMP1/MMP1/MMP3 | 3 |
| GO:0034381 | plasma lipoprotein particle clearance | 0.000744 | 0.017445 | 0.014601 | APOM/APOA1/APOE | 5 |
| GO:0002526 | acute inflammatory response | 0.000746 | 0.017445 | 0.014601 | F2/A2M/C3/TFRC/SER | 7 |
| GO:0031424 | keratinization | 0.000759 | 0.017638 | 0.014763 | KRT16/KRT5/CASP14 | 6 |
| GO:0010743 | regulation of macrophage derived foam cell differentiation | 0.000764 | 0.017638 | 0.014763 | AGT/APOB/ITGAV/C | 4 |
| GO:0030324 | lung development | 0.000775 | 0.017761 | 0.014866 | LAMA5/RBP4/LAMA | 9 |
| GO:0007178 | transmembrane receptor protein serine/threonine kinase si | 0.00078 | 0.017761 | 0.014866 | GREM1/AFP/TF/HEX | 14 |
| GO:0051222 | positive regulation of protein transport | 0.000787 | 0.017809 | 0.014906 | GLUD1/F2/FGB/RBP4 | 12 |
| GO:0002253 | activation of immune response | 0.00083 | 0.018656 | 0.015615 | CFD/A2M/C3/C5/C9/L | 16 |
| GO:0061138 | morphogenesis of a branching epithelium | 0.000836 | 0.018671 | 0.015628 | NRP1/LAMA5/GREM | 9 |
| GO:0001667 | ameboidal-type cell migration | 0.000848 | 0.018812 | 0.015746 | NRP1/LAMA5/GREM | 16 |
| GO:0006826 | iron ion transport | 0.000876 | 0.01932 | 0.016171 | TFRC/TF/LTF/FTL/FT | 5 |

| | | | | | | |
|---|---|---|---|---|---|---|
| GO:0030323 | respiratory tube development | 0.000901 | 0.019742 | 0.016524 | LAMA5/RBP4/LAMA | 9 |
| GO:0032092 | positive regulation of protein binding | 0.000973 | 0.021048 | 0.017617 | NRP1/APOE/VTN/RPL | 6 |
| GO:0097006 | regulation of plasma lipoprotein particle levels | 0.000973 | 0.021048 | 0.017617 | APOM/AGT/APOA1/A | 6 |
| GO:0002719 | negative regulation of cytokine production involved in imr | 0.00108 | 0.022626 | 0.018938 | APOA1/AXL/ANGPT | 4 |
| GO:0030214 | hyaluronan catabolic process | 0.001085 | 0.022626 | 0.018938 | HEXA/HEXB/CEMIP | 3 |
| GO:0034374 | low-density lipoprotein particle remodeling | 0.001085 | 0.022626 | 0.018938 | AGT/APOE/APOB | 3 |
| GO:0034375 | high-density lipoprotein particle remodeling | 0.001085 | 0.022626 | 0.018938 | APOM/APOA1/APOE | 3 |
| GO:0071732 | cellular response to nitric oxide | 0.001085 | 0.022626 | 0.018938 | MMP3/ATP5F1A/HNR | 3 |
| GO:0090136 | epithelial cell-cell adhesion | 0.001085 | 0.022626 | 0.018938 | JUP/DSP/VCL | 3 |
| GO:0002237 | response to molecule of bacterial origin | 0.001094 | 0.022665 | 0.018971 | SSC5D/PENK/LTF/AP | 13 |
| GO:0002698 | negative regulation of immune effector process | 0.001112 | 0.022916 | 0.01918 | A2M/HLA-B/APOA1/A | 7 |
| GO:0031647 | regulation of protein stability | 0.001123 | 0.023002 | 0.019252 | APOA1/TF/GAPDH/C | 12 |
| GO:0033687 | osteoblast proliferation | 0.001202 | 0.024474 | 0.020485 | GREM1/LTF/ITGAV/C | 4 |
| GO:0010631 | epithelial cell migration | 0.001236 | 0.025012 | 0.020935 | NRP1/GREM1/AGT/A | 13 |
| GO:0032488 | Cdc42 protein signal transduction | 0.001305 | 0.026093 | 0.021839 | NRP1/APOA1/APOE | 3 |
| GO:1903829 | positive regulation of protein localization | 0.001305 | 0.026093 | 0.021839 | GLUD1/F2/FGB/RBP4 | 15 |
| GO:0090132 | epithelium migration | 0.001328 | 0.026093 | 0.021839 | NRP1/GREM1/AGT/A | 13 |
| GO:0044273 | sulfur compound catabolic process | 0.001334 | 0.026093 | 0.021839 | HEXA/HEXB/GPC1/N | 4 |
| GO:0043393 | regulation of protein binding | 0.001337 | 0.026093 | 0.021839 | NRP1/APOE/VTN/LO | 9 |
| GO:0002688 | regulation of leukocyte chemotaxis | 0.001342 | 0.026093 | 0.021839 | ADAM10/GREM1/C5/ | 7 |
| GO:0003206 | cardiac chamber morphogenesis | 0.001342 | 0.026093 | 0.021839 | NRP1/RBP4/DSP/CPE/ | 7 |
| GO:0008544 | epidermis development | 0.00136 | 0.026303 | 0.022015 | LAMA5/KRT16/KRT5/ | 13 |
| GO:0031638 | zymogen activation | 0.001381 | 0.026554 | 0.022225 | FGB/ENO1/SERPINF2 | 5 |
| GO:0001763 | morphogenesis of a branching structure | 0.001431 | 0.027371 | 0.02291 | NRP1/LAMA5/GREM | 9 |
| GO:1990266 | neutrophil migration | 0.00147 | 0.027915 | 0.023365 | LBP/ITGA1/PPIA/TNF | 7 |
| GO:0010742 | macrophage derived foam cell differentiation | 0.001476 | 0.027915 | 0.023365 | AGT/APOB/ITGAV/C | 4 |
| GO:0090130 | tissue migration | 0.001495 | 0.028124 | 0.02354 | NRP1/GREM1/AGT/A | 13 |
| GO:1903532 | positive regulation of secretion by cell | 0.001516 | 0.028367 | 0.023743 | GLUD1/F2/AGT/FGB/ | 11 |
| GO:0006029 | proteoglycan metabolic process | 0.001536 | 0.028555 | 0.023901 | HEXA/HEXB/BGN/G | 6 |
| GO:0034384 | high-density lipoprotein particle clearance | 0.001551 | 0.028555 | 0.023901 | APOM/APOA1/APOE | 3 |
| GO:1902170 | cellular response to reactive nitrogen species | 0.001551 | 0.028555 | 0.023901 | MMP3/ATP5F1A/HNR | 3 |
| GO:0071692 | protein localization to extracellular region | 0.001603 | 0.02933 | 0.024549 | GLUD1/F2/APOE/FG | 13 |

| | | | | | | |
|---|---|---|---|---|---|---|
| GO:0052548 | regulation of endopeptidase activity | 0.00161 | 0.02933 | 0.024549 | A2M/TIMP1/LTF/VTN | 12 |
| GO:0090077 | foam cell differentiation | 0.001628 | 0.029501 | 0.024692 | AGT/APOB/ITGAV/CS | 4 |
| GO:0002697 | regulation of immune effector process | 0.001679 | 0.030269 | 0.025335 | A2M/C3/HLA-B/APOA | 13 |
| GO:0016485 | protein processing | 0.00171 | 0.030673 | 0.025673 | ADAM10/FGB/ENO1/ | 10 |
| GO:0010744 | positive regulation of macrophage derived foam cell differ | 0.001824 | 0.032379 | 0.027101 | AGT/APOB/CSF1 | 3 |
| GO:0030540 | female genitalia development | 0.001824 | 0.032379 | 0.027101 | RBP4/AXL/LRP2 | 3 |
| GO:0060541 | respiratory system development | 0.001866 | 0.032959 | 0.027586 | LAMA5/RBP4/LAMA | 9 |
| GO:0098869 | cellular oxidant detoxification | 0.001897 | 0.03333 | 0.027897 | APOM/HBE1/APOE/A | 6 |
| GO:0032496 | response to lipopolysaccharide | 0.002007 | 0.035076 | 0.029358 | PENK/LTF/APOB/CSF | 12 |
| GO:0045216 | cell-cell junction organization | 0.002055 | 0.035607 | 0.029803 | ADAM10/AGT/RHOC | 9 |
| GO:0050900 | leukocyte migration | 0.002058 | 0.035607 | 0.029803 | ADAM10/GREM1/C5/ | 13 |
| GO:0030216 | keratinocyte differentiation | 0.002078 | 0.035786 | 0.029953 | KRT16/KRT5/DSP/CA | 8 |
| GO:0050764 | regulation of phagocytosis | 0.0021 | 0.035888 | 0.030038 | C3/APOA1/ITGAV/TG | 6 |
| GO:0043691 | reverse cholesterol transport | 0.002126 | 0.035888 | 0.030038 | APOM/APOA1/APOE | 3 |
| GO:0044794 | positive regulation by host of viral process | 0.002126 | 0.035888 | 0.030038 | APOE/STOM/EEF1A1 | 3 |
| GO:0046514 | ceramide catabolic process | 0.002126 | 0.035888 | 0.030038 | HEXA/HEXB/ASAH1 | 3 |
| GO:0045927 | positive regulation of growth | 0.002154 | 0.036195 | 0.030295 | AGRN/ADAM10/NRP | 10 |
| GO:0048511 | rhythmic process | 0.002197 | 0.036581 | 0.030618 | A2M/AFP/AXL/PTGD | 11 |
| GO:0030316 | osteoclast differentiation | 0.002207 | 0.036581 | 0.030618 | TFRC/TF/LTF/CSF1/FI | 6 |
| GO:0050922 | negative regulation of chemotaxis | 0.002209 | 0.036581 | 0.030618 | NRP1/GREM1/C5/TNF | 5 |
| GO:2001235 | positive regulation of apoptotic signaling pathway | 0.002449 | 0.040289 | 0.033722 | AGT/CTSC/RPS7/RPL | 7 |
| GO:0071731 | response to nitric oxide | 0.002456 | 0.040289 | 0.033722 | MMP3/ATP5F1A/HNR | 3 |
| GO:0006024 | glycosaminoglycan biosynthetic process | 0.002501 | 0.040823 | 0.034169 | B4GAT1/HEXA/ANGI | 5 |
| GO:1900026 | positive regulation of substrate adhesion-dependent cell sp | 0.002554 | 0.041496 | 0.034732 | NRP1/APOA1/FGB/M7 | 4 |
| GO:0030449 | regulation of complement activation | 0.002817 | 0.044756 | 0.037461 | A2M/C3/SVEP1 | 3 |
| GO:0035089 | establishment of apical/basal cell polarity | 0.002817 | 0.044756 | 0.037461 | LAMA1/FAT1/PATJ | 3 |
| GO:0051444 | negative regulation of ubiquitin-protein transferase activity | 0.002817 | 0.044756 | 0.037461 | RPS7/RPL11/RPL5 | 3 |
| GO:2001044 | regulation of integrin-mediated signaling pathway | 0.002817 | 0.044756 | 0.037461 | TIMP1/LAMA1/LAME | 3 |
| GO:0006023 | aminoglycan biosynthetic process | 0.002819 | 0.044756 | 0.037461 | B4GAT1/HEXA/ANGI | 5 |
| GO:0042742 | defense response to bacterium | 0.002891 | 0.045692 | 0.038244 | SSC5D/F2/FGB/TF/LT | 11 |
| GO:0042176 | regulation of protein catabolic process | 0.002908 | 0.045746 | 0.038289 | SEC22B/TIMP1/APOE | 12 |
| GO:0044788 | modulation by host of viral process | 0.003008 | 0.04711 | 0.039431 | APOE/LTF/STOM/EEF | 4 |

| | | | | | | |
|---|---|---|---|---|---|---|
| GO:0050714 | positive regulation of protein secretion | 0.003097 | 0.048242 | 0.040378 | GLUD1/F2/FGB/RBP4 | 7 |
| GO:0051047 | positive regulation of secretion | 0.003108 | 0.048242 | 0.040378 | GLUD1/F2/AGT/FGB/ | 11 |
| GO:0002689 | negative regulation of leukocyte chemotaxis | 0.003208 | 0.049489 | 0.041422 | GREM1/C5/TNFAIP6 | 3 |
| GO:0002701 | negative regulation of production of molecular mediator of | 0.003255 | 0.049489 | 0.041422 | APOA1/AXL/ANGPT1 | 4 |
| GO:0043277 | apoptotic cell clearance | 0.003255 | 0.049489 | 0.041422 | C3/ITGAV/TGM2/AXI | 4 |
| GO:0061383 | trabecula morphogenesis | 0.003255 | 0.049489 | 0.041422 | GREM1/RBP4/FBN2/A | 4 |
| GO:0006790 | sulfur compound metabolic process | 0.00326 | 0.049489 | 0.041422 | PHGDH/B4GAT1/HEX | 11 |

**Supplementary Table 2: Comparison of data with published data [31]**

**Overexpressed proteins**

| SL No | UniProt Accession | Description of the protein | Entrez Gene ID | Ensembl Gene ID | Gene Symbol | log2FC from our study | p-value from our study | log2FC from available study | p-value from available study |
|---|---|---|---|---|---|---|---|---|---|
| 1 | O60565 | Gremlin-1 | 26585 | ENSG00000166923; ENSG00000276886; ENSG00000282046 | GREM1 | 3.124969244 | 0.00482142 | 3.09 | 0.01 |

**Under-expressed proteins**

| SL No | UniProt Accession | Description of the protein | Entrez Gene ID | Ensembl Gene ID | Gene Symbol | log2FC from our study | p-value from our study | log2FC from available study | p-value from available study |
|---|---|---|---|---|---|---|---|---|---|
| 1 | P08254 | Stromelysin-1 | 4314 | ENSG00000149968 | MMP3 | -4.428064395 | 0.00055209 | -2.29 | 0.002 |
| 2 | P41222 | Prostaglandin-H2 D-isomeras | 5730 | ENSG00000107317 | PTGDS | -1.601496461 | 0.014555 | -3.38 | 0.019 |
| 3 | P98095 | Fibulin-2 | 2199 | ENSG00000163520 | FBLN2 | -3.863706093 | 0.00883762 | -3.4 | 0.012 |
| 4 | P02649 | Apolipoprotein E | 348 | ENSG00000130203 | APOE | -3.266212818 | 0.00064402 | -4.45 | 0.003 |
